# Static Kinks or Flexible Hinges: Conformational Distributions of Bent DNA Bound to Integration Host Factor Mapped by Fluorescence Lifetime Measurements

**DOI:** 10.1101/382655

**Authors:** Mitchell Connolly, Aline Arra, Viktoriya Zvoda, Peter J. Steinbach, Phoebe A. Rice, Anjum Ansari

**Affiliations:** Department of Physics, University of Illinois at Chicago, Chicago, IL 60607, USA; Center for Molecular Modeling, Center for Information Technology, National Institutes of Health, Bethesda, MD 20892, USA; Department of Biochemistry & Molecular Biology, University of Chicago, Chicago, IL 60637,USA; Department of Bioengineering, University of Illinois at Chicago, Chicago, IL 60607, USA

## Abstract

Gene regulation depends on proteins that bind to specific DNA sites. Such specific recognition often involves severe DNA deformations including sharp kinks. It has been unclear how rigid or flexible these protein-induced kinks are. Here, we investigated the dynamic nature of DNA in complex with integration host factor (IHF), a nucleoid-associated architectural protein known to bend one of its cognate sites (35 base pair H’) into a U-turn by kinking DNA at two sites. We utilized fluorescence lifetime based FRET spectroscopy to map the distribution of bent conformations in various IHF-DNA complexes. Our results reveal a surprisingly dynamic specific complex: while 80% of the IHF-H’ population exhibited FRET efficiency consistent with the crystal structure, 20% exhibited FRET efficiency indicative of unbent or partially bent DNA. This conformational flexibility is modulated by sequence variations in the cognate site. In another site (H1) that lacks an A-tract of H’ on one side of the binding site, the population in the fully U-bent conformation decreased to 36%, as did the extent of bending. A similar decrease in the U-bent population was observed with a single base mutation in H’ in a consensus region on the other side. Taken together, these results provide important insights into the finely tuned interactions between IHF and its cognate sites that keep the DNA bent (or not), and yield quantitative data on the dynamic equilibrium between different DNA conformations (kinked or not kinked) that depend sensitively on DNA sequence and deformability. Notably, the difference in dynamics between IHF-H’ and IHF-H1 reflects the different roles of these complexes in their natural context, in the phage lambda “intasome” (the complex that integrates phage lambda into the *E. coli* chromosome).

## INTRODUCTION

Severe distortions to the B-DNA structure are ubiquitous in the cell and are induced not only by nonspecific DNA packaging proteins but also by site-specific proteins when they bind to their target sites on DNA for gene regulation, replication and repair. These distortions often appear as DNA bends or localized kinks that range from ˜30° kinks per helical turn when wrapped in the nucleosome,^1–4^ to more severe kinks of ˜80-90°in several site-specific DNA bending proteins,^5–8^ to what may be the most severely bent DNA conformation found in complex with the IHF/HU family of bacterial type II DNA-bending proteins.^9–11^ While structural studies on many DNA-bending protein complexes have revealed the nature of these DNA distortions at atomic resolution, the behavior of the distorted, sharply kinked, DNA in solution needs further study. One question that has yet to be fully resolved is whether these distorted DNA structures behave as rigid kinks in solution or as flexible hinges.^12^ Precise measurements of the accessible protein-DNA conformations in solution are needed to elucidate the functional roles of these dynamic fluctuations as well as to provide experimental checkpoints in the development and improvement of computational models of sequence-dependent DNA deformability and stabilization of bent DNA by proteins.^13–22^

Several lines of evidence suggest that DNA in nonspecific complexes indeed adopts a range of bent conformations. Evidence for conformational heterogeneity is well documented in the case of the nonspecific architectural histone-like nucleoprotein HU that is known to bend DNA into a U-shape.^10, 23^ Indirect evidence for multiple bent states in HU-DNA complexes came from multiple bands revealed in gel shift assays,^24^ and from different degrees of bending observed in different crystal structures of these complexes.^10^ This heterogeneity in HU-DNA complexes was directly confirmed from atomic force microscopy (AFM) studies that showed a broad distribution of bent conformations with bend angles observed in the entire range from 0 to 180°.^25^ It is perhaps not too surprising that a nonspecific architectural DNA bending protein, especially one that has a variety of biological roles, would exhibit such broad conformational heterogeneity, given the lack of specific contacts that would otherwise compensate for the energetic penalty required to severely deform DNA.

However, are specific complexes with DNA bending proteins also conformationally heterogeneous? AFM studies on some specific complexes have revealed relatively broad distributions of bend angles, although the resolution of these AFM studies primarily revealed single-peaked distributions of varying widths, indicative of thermal fluctuations within a single free energy well in the conformational landscape of the complex.^26–28^ Notable exceptions in which two or more distinct bent states of DNA are observed include mismatch repair protein MutS^29^ and DNA glycosylase hTDG^30^ bound to mismatched DNA. Interestingly, these are examples of DNA damage recognition proteins in which the specific site is typically a single mismatch or a single chemically modified base, and all other interactions of the protein with the flanking DNA sequences are nonspecific. For site-specific proteins that bind to canonical, undamaged DNA, and for which the target site extends beyond a few base pairs, data are scarce regarding conformational heterogeneity in such complexes.

Here, we present results that quantify the heterogeneity of bent DNA conformations when bound to the eukaryotic DNA-bending protein, the *Escherichia coli* integration host factor (IHF). We utilize picosecond-resolved fluorescence lifetime measurements to map the distribution of FRET states, which reflects the distribution of bent conformations in the IHF-DNA complex. The lifetime approach has a few advantages over single-molecule assays of FRET distributions.^31^ Measurements are done under solution conditions and thus avoid many of the issues that may come with immobilization of protein-DNA complexes as needed for AFM studies^32–33^ or single-molecule FRET (smFRET) studies,^34–37^ although smFRET on freely diffusing molecules do alleviate immobilization issues, as showcased in a series of elegant papers by Eaton and co-workers^38–41^ on the dynamics and distributions during protein folding. More important, fluorescence decay curves measured in bulk solution provide significantly better resolution of the shape of the FRET efficiency distributions than can be obtained from the few-to-tens thousands molecules that are sampled in single-molecule studies. Moreover, the time resolution of the lifetime studies enables snapshots of the conformational distribution with an effective “shutter speed” that is typically less than tens of nanoseconds (within the excited state lifetime of the fluorophore), compared with few milliseconds or longer in smFRET, which can obscure conformational heterogeneity if dynamic fluctuations between different conformations are faster than a few milliseconds.^42^ Recently, we demonstrated the effectiveness of fluorescence lifetime based FRET measurements in unveiling the conformational heterogeneity in mismatched DNA bound to nucleotide excision repair protein XPC/Rad4, with evidence for two or more distinct DNA conformations in the complex.^43^

IHF is a small (22 kDa) bacterial heterodimeric nucleoid-associated protein that is closely related to HU and structurally very similar.^23, 44^ Unlike HU, however, IHF binds DNA both nonspecifically, in its role as a DNA compaction protein, and specifically, when required for site-specific recombination, DNA replication and transcription. IHF also recognizes several sites on bacteriophage lambda DNA in its role as a host factor for lysogeny by phage lambda.^45–46^ The footprint of DNA bound to IHF exceeds 25 base pairs (bp).^47^ However, the consensus sequence identified on the basis of a large number of cognate sites for IHF consists of two short elements in one half of the footprinted region (Figure 1): WATCARnnnnTTR, where W denotes A or T, R denotes purine, and n refers to any base.^47–48^ Some IHF sites also contain an A-tract containing 4-6 adenines and located in the other half of the binding site.^49–50^

**Figure 1.**
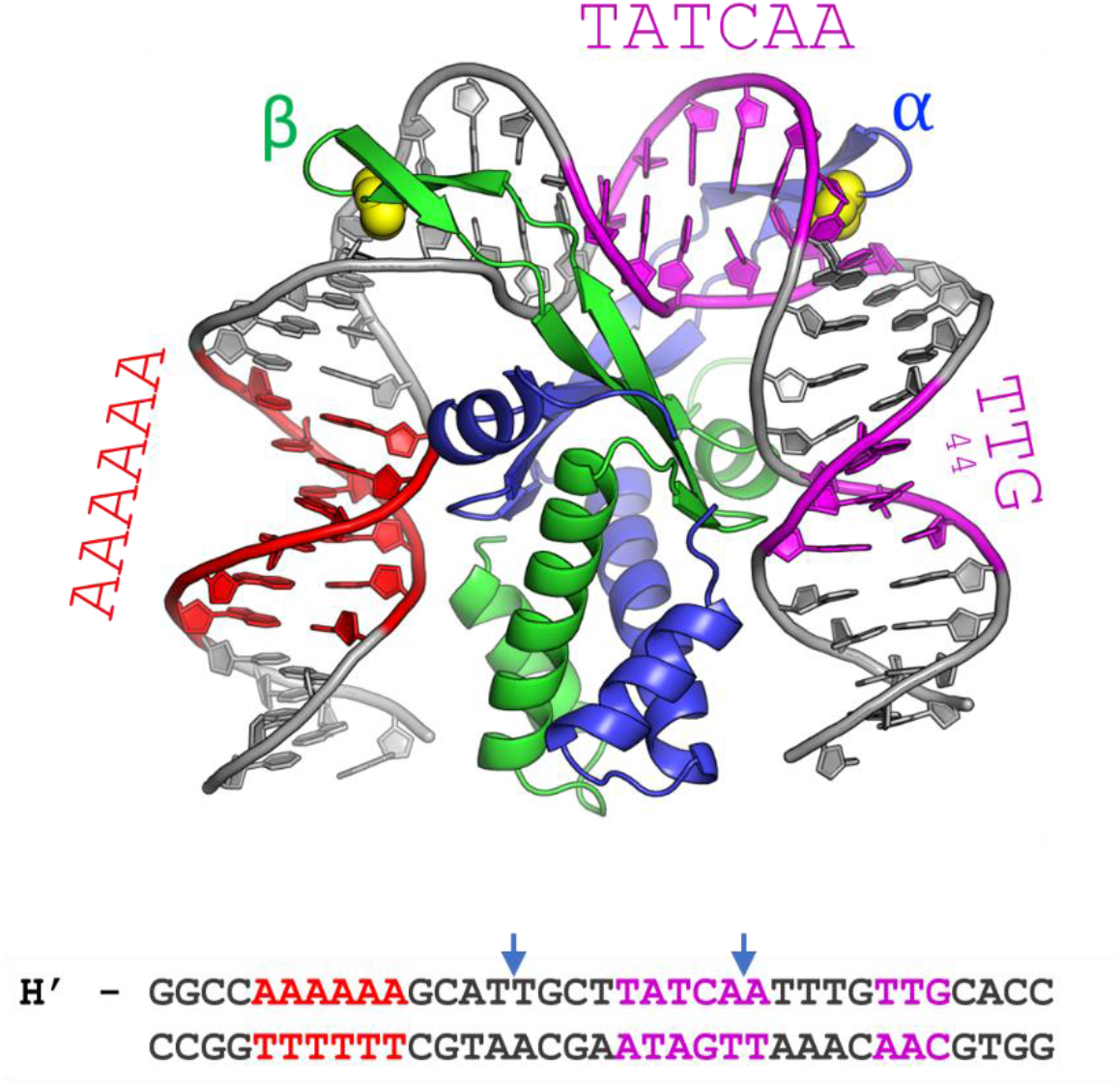
The cocrystal structure of IHF in complex with the H’ binding site from phage lambda. The α- and β-chains of the IHF protein are shown in blue and green, respectively, with the conserved proline residues shown as yellow spheres. The DNA is shown in gray, with the consensus region highlighted in magenta and the A-tract in red. The sequence shown below is that of the 35-mer that contains the H’ binding site, which was used to obtain the cocrystal structure with IHF. In the complex, the DNA is sharply kinked at the two sites indicated by the blue arrows. In the DNA oligomer used for the structural studies, the DNA was nicked at a position shifted 1 bp to the 3’-side of the left blue arrow, to facilitate crystal packing. That nick was “sealed” *in silico* to generate the model shown (starting from PDB ID 1IHF). Figure made using the PyMol molecular graphics system, version 2.0, Schrodinger, LLC.

The crystal structure of IHF bound to one such cognate site, denoted as the H’ site (Figure 1 and Table 1), showed that IHF sharply kinks the DNA at two sites spaced ˜9 bp apart, explaining how it can bring distal regions of DNA together to facilitate the formation of higher-order nucleoprotein complexes.^9^ The kinks in the DNA are stabilized by conserved proline residues located on two β-ribbon arms of the protein. The arms are thought to be flexible in the absence of the DNA but wrap around the DNA in the complex and make additional stabilizing contacts in the consensus region between the kink sites (Figure 1). Further stabilization needed to overcome the large energy penalty for the sharply kinked DNA comes from additional contacts between the flanking DNA segments and the core of the protein dimer, as well as from an extensive network of electrostatic interactions with charged residues of the protein and the phosphates in the DNA backbone.^51–52^

**Table 1:**
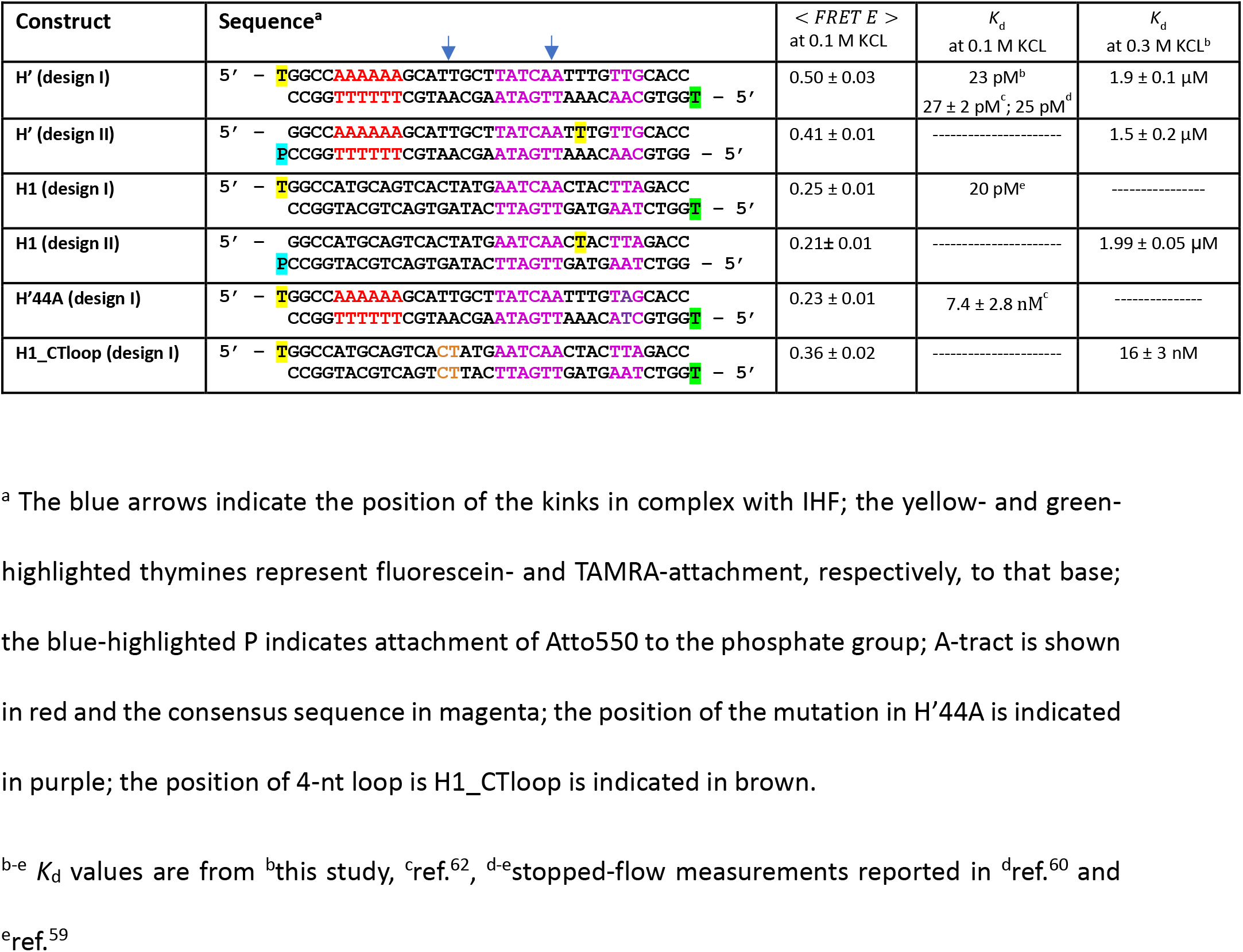
DNA Constructs Used in This Study

All contacts that IHF makes with the DNA in the specific complex are either in the minor groove, where the hydrogen bonding patterns offered by the different bases are very similar,^53^ or to the sugar phosphate backbone. Thus, IHF is a remarkable example of a DNA-bending protein that exhibits high specificity to certain sequences but relies almost exclusively on “indirect readout” to recognize its target sequence; i.e. it recognizes sequence-dependent DNA shape, local geometry, and DNA deformability without the need for direct recognition of the specific bases by the protein residues.^54^ Several features within the binding site play a role in facilitating this recognition. One may be the ease with which the two sites can be kinked: although only one kink occurs within the consensus sequence, introducing single-T insertions or mismatches to make the DNA more flexible at these sites was shown to increase the binding affinity for IHF as well as for the nonspecific HU.^55–56^ Another important feature is the flanking DNA sequences that interact with the sides of the protein, where the minor groove of the DNA is clamped between the N-termini of two alpha helices. On the right side (as shown in Figure 1 and Table 1), the TTR of the consensus sequence allows over-twisting of the DNA that facilitates its fit into the protein clamp and the formation of salt bridges with the side chains of IHF.^57^ On the left side of H’ (but not all IHF binding sites) is an A-tract. The structure shows that the unusually narrow groove naturally adopted by A-tract DNA fits well into the protein clamp, and this may explain why crystals could not be grown with IHF binding sites that did not include the A-tract.^9^ This study seeks to elucidate how each of these features help keep the DNA bent and clamped against the protein and to what extent the DNA resists these bending deformations.

Here, we present fluorescence lifetime-based FRET measurements to demonstrate that IHF bound to a 35-bp substrate containing the cognate H’ site, the sequence used in the crystal structure, in fact samples two distinct conformations: one that appears to be fully bent, as in the crystal structure, and another that appears to be only partially bent and could be an ensemble of conformations with one or the other side unclamped. Another cognate site that is lacking the A-tract (the H1 site on phage lambda DNA) showed a significantly smaller population in the fully bent state, highlighting the effectiveness of the A-tract to keep that side of the DNA clamped down. The population in the fully bent state in the IHF-H1 complex is partially recovered when mismatches are introduced at the kink site near the missing A-tract, to reduce the energetic cost of kinking the DNA at that site. Finally, a single base modification in the TTR consensus region in the other flanking arm, previously known to destabilize the complex, also resulted in a significantly diminished population in the fully bent state. Taken together, these results provide additional insights into the finely tuned interactions between IHF and its cognate sites that keep the DNA bent (or not) and yield quantitative data on the dynamic fluctuations between different DNA conformations (kinked or not kinked) that are sensitively dependent on the DNA sequence and deformability. These results also correlate well with the biological function of the two IHF-binding sites studied. These two sites are found within the large protein-DNA complex responsible for integrating phage lambda DNA into the *E. coli* chromosome, and structural modeling shows that the IHF-induced bend at the H1 site, but not at the bend H’ site, must flex during assembly.^58^

## EXPERIMENTAL METHODS

### Materials

The DNA sequences used in this study are shown in Table 1. All labeled and unlabeled DNA oligomers were ordered from Keck (with gel purification) or from IDT (with HPLC purification). Two different labeling strategies were used as shown in Table 1: in design I constructs, fluorescein (F) and TAMRA (R) were attached to thymidine overhangs at the at 5′-end of the top and bottom strands, respectively, through six-carbon phosphoradimite linkers; in design II constructs, fluorescein (F) was attached to a thymidine located 10 nucleotides from the 3′-end of the top strand and two base pairs away from one of the kink sites, and Atto550 (At) was attached to the bottom strand, to the phosphate group of the DNA backbone at the 3′ end, also by six-carbon phosphoradimite linkers.

DNA concentrations were determined by absorbance measurements at 260nm, with extinction coefficients 3.58×10^5^ M^−1^cm^−1^ for the 5′-end F-labeled (top) strand, 3.66×10^5^ M^−1^cm^−1^ for the 5′-end R-labeled (bottom) strand, 3.48×10^5^ M^−1^cm^−1^ for the mid-F-labeled (top) strand and 3.68×10^5^ M^−1^cm^−1^ for the 3′-end At-labeled (bottom) strand. Labeling efficiencies were determined by simultaneous absorbance measurements on F-labeled strands at 494 nm (molar extinction coefficient 75,000 M^−1^cm^−1^); on R-labeled strands at 555 nm (molar extinction coefficient 91,000 M^−1^cm^−1^); and on At-stands at 560 nm (molar extinction coefficient 131,200 M^−1^cm^−1^); extinction coefficients were obtained from Molecular Probes http://www.glenresearch.com/Technical/Extinctions.html). The percentage of labeled DNA in solution was estimated to be >87% for all donor-labeled strands and >95% for all acceptor-labeled strands.

Duplex DNA was formed by annealing complimentary oligomers of equimolar concentrations. The annealing buffer used was 20 mM Tris-HCl, pH 8.0, 1 mM ethylenediaminetetraacetic acid (EDTA), and 200 mM KCl. The mixture of oligomers was heated in a water bath at 90 °C for 10 minutes, then allowed to cool slowly at room temperature.

The IHF protein was prepared as described previously (64). Droplets of proteins were first flash-frozen in liquid nitrogen prior to storage in cryogenic tubes at −80 °C. Individual frozen droplets were diluted into the binding buffer, as needed. All measurements were performed in binding buffer: 20 mM Tris–HCl (pH 8.0), 1 mM EDTA, 0.01% NP-40 with salt concentrations ranging from 100 to 300 mM KCl. Protein concentrations reported in this study are effectively “active” protein concentrations, determined as described in SI Methods 1.1.

### Steady-state fluorescence (acceptor ratio and anisotropy) measurements

The steady-state fluorescence emission spectra and anisotropies were measured on a FluoroMax4 spectrofluorometer (Jobin Yvon, Inc., NJ, USA), with samples loaded in a 100-μL quartz cuvette (Starna 26.100F-Q-10/Z20). Details of the acceptor ratio and anisotropy measurements are in SI Methods 1.2.

### Binding affinity measurements

To determine the effect of the different labeled sequences on the IHF binding affinities, we performed equilibrium titration measurements for IHF and different DNA constructs at 20 °C. These comparative binding affinity measurements were done at 300 mM KCl, to bring the dissociation constants of the complexes (*K_d_*) in the > nM range. At 100 mM KCl, where most of the lifetime measurements were done, the *K_d_* for the IHF-H’ complex was previously found to be in the pM range,^59–62^ such that conventional titration experiments cannot measure these *K_d_* values accurately.^61^ Details of the binding affinity measurements are in SI Methods 1.3.

### Fluorescence lifetime spectroscopy

Fluorescence decay curves were measured with a PicoMaster fluorescence lifetime spectrometer (HORIBA-PTI, London, Ontario, Canada) equipped with time-correlated single photon counting (TSCPC) electronics.^63^ For all FRET measurements, decay traces were measured for donor-only duplexes without acceptor, denoted as DNA_D, as well as donor-acceptor-labeled duplexes, denoted as DNA_DA. The excitation source was a Fianium Whitelase Supercontinuum laser system (maximum power 4W), which produces ˜6 ps broad band pulses. For excitation of F, the laser pulses were passed through a monochromator set at 485 nm (bandpass 10 nm) followed by a 488 ± 10 nm bandpass filter. The emission from the sample was collected orthogonal to the excitation beam after passing through a 496 nm longpass filter (Semrock BrightLine FF01-496/LP-25), followed by another monochromator set at 520 nm (bandpass 10 nm), and detected by a Hamamatsu microchannel plate photomultiplier (MCP-650). The instrument response function (IRF) of the system was measured using a dilute aqueous solution of Ludox (Sigma-Aldrich). The full width at half maximum (fwhm) of the IRF was ˜100 ps. Fluorescence decay curves were recorded on a 100 ns timescale, resolved into 4096 channels, to a total of 10,000 counts in the peak channel, with the repetition rate of the laser adjusted to 10 MHz.

### Maximum entropy analysis of the fluorescence decay traces

Fluorescence decay curves were analyzed using a maximum entropy method (MEM) in which the effective distribution of log-lifetimes *f*(log τ) was inferred from the decay traces using the program MemExp (available online), and described in detail elsewhere.^64–65^ The signal measured at time *t_i_* was fit by the expression:

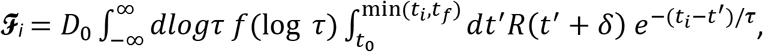

where *R* is the measured instrument response function, δ is the zero-time shift and *D_0_* is a normalization constant. The instrument response is appreciable in the range [*t*_0_, *t_f_*]. The zerotime shift was determined using Brent’s method of optimization and preliminary MEM calculations, each performed with δ fixed at a different value. A similar estimation of δ has been reported.^66^ The Poisson deviance between the fit and data was minimized while maximizing the entropy of the *f* distribution.

The MEM inverts fluorescence decay traces into lifetime distributions without any *a priori* assumptions about the number of exponential terms. The MEM outputs of the donor-only samples gave single-peaked, narrow distributions, consistent with single-exponential decay. Fits to discrete exponential decays with two or more exponentials yielded less than ˜1% amplitude in the additional decay components (SI Figure S1).

The average FRET efficiencies measured on donor-acceptor labeled (DNA_DA) samples were computed from the MEM distributions as described in SI Methods 1.4. In the presence of the protein, lifetime decays on some DNA_DA samples yielded bimodal distributions, which were further analyzed in terms of two components, as described in SI Methods 1.4.

## RESULTS AND DISCUSSION

### Fluorescence lifetime measurements provide a snapshot of the distribution of bent conformations in IHF-DNA complexes

In this study, we mapped the equilibrium distribution of DNA conformations when bound to IHF, by measuring the FRET efficiency between a donor-acceptor FRET pair attached to a 35-bp DNA substrate of varying sequences and binding affinities, using fluorescence lifetime studies. The FRET efficiency (E) between the donor and acceptor labels depends strongly on the relative distance and orientation between the labels and can be computed from the lifetimes of the excited donor fluorophore in the presence (*τ_DA_*) and absence (*τ_D_*) of the acceptor, as 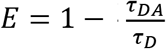. The measured FRET efficiencies can sense changes in the conformations of the DNA to which the labels are attached, for example upon binding of a DNA-bending protein. In the case of IHF-DNA complexes, FRET between labels attached at the ends of 35-mer DNA substrates has proven to be exceptionally useful in measuring not only equilibrium changes from unbent (straight) to bent conformations upon binding of IHF,^67^ but also the dynamics of these conformational changes along the recognition trajectory of the IHF binding site.^62, 68–69^

Here, we take advantage of the sub-nanosecond time-resolution of the fluorescence lifetime studies that enables fluorescence decays to be measured over a wide temporal range and with high temporal resolution. A single DNA conformation in the ensemble of molecules, with a well-defined separation and relative orientation between the donor and acceptor of the FRET pair, is characterized by a unique FRET efficiency, and is expected to yield a single-exponential decay profile in the lifetime measurements. A distribution of DNA conformations corresponding to different donor-acceptor distance/orientations would then correspond to a distribution of FRET efficiencies, which should be reflected in a distribution of lifetimes measured in the decay traces. The lifetime distributions were obtained from the measured decay traces by the maximum entropy method (MEM).

The fluorescence lifetime of the donor (fluorescein), measured in DNA constructs in the absence of any acceptor (DNA_D), exhibited close to single-exponential decays with a relatively narrow distribution of lifetimes, and characterized by an effectively unique lifetime < *τ_D_* > ≈ 4.2 – 4.3 ns that was found to be mostly insensitive to the position of attachment to the DNA or to the presence of the protein (SI Figure S1). In the presence of an acceptor fluorophore (DNA_DA), the decay profiles remained largely single-exponential with no protein bound (Figure 2), albeit with a shift in the donor lifetime < *τ_DA_* > relative to < *τ_D_* > if there was significant FRET (SI Figure S2). In contrast, for lifetime measurements on DNA_DA in the presence of IHF, not only did the donor lifetime shorten due to FRET, we also detected a broadening of the distribution of lifetimes and appearance of at least two distinct components (Figure 2 and SI Figure S3), which points to direct evidence for multiple DNA conformations in the IHF-bound complexes in the ensemble.

**Figure 2.**
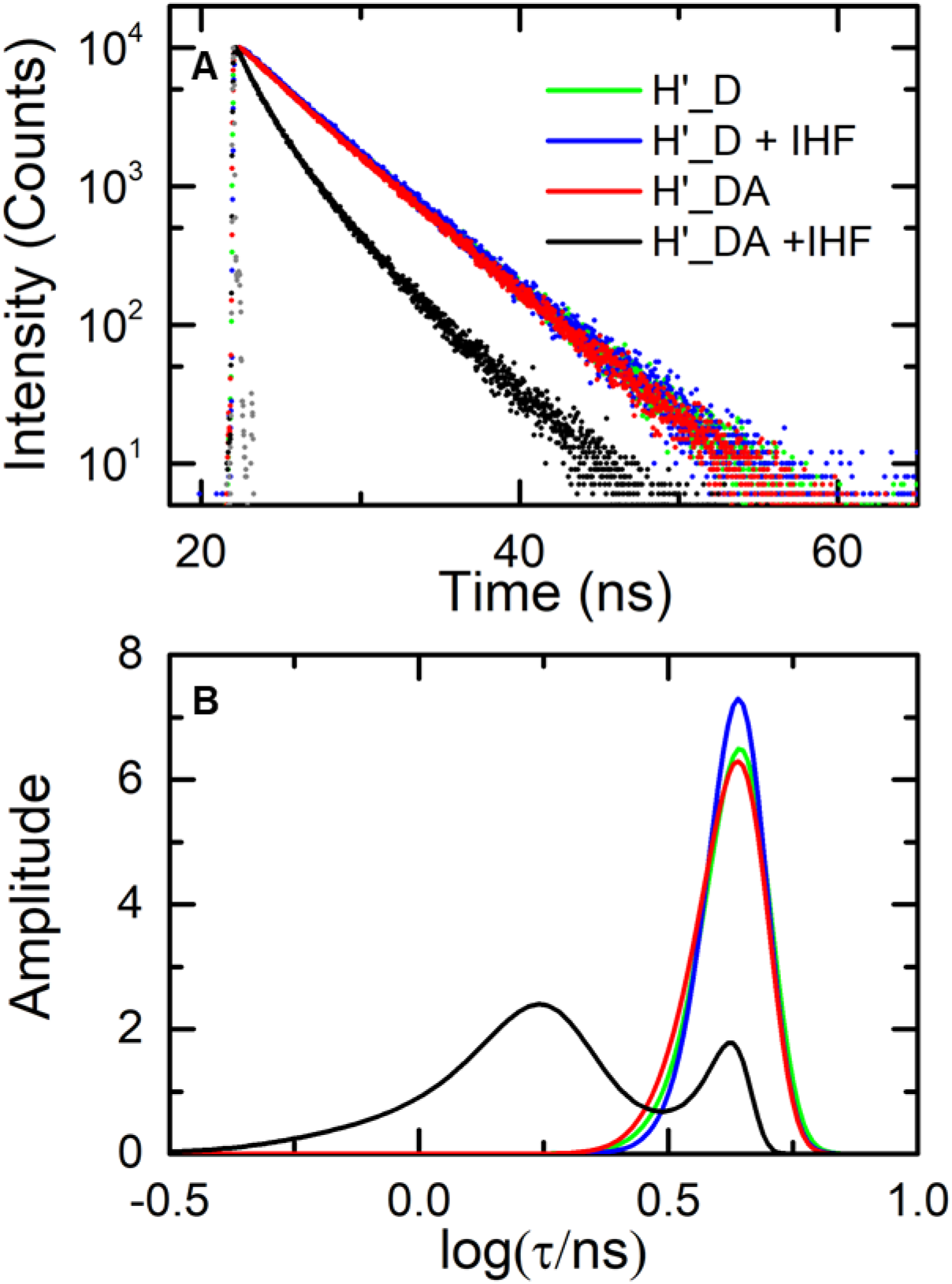
Fluorescence lifetime measurements on IHF-H’ complex in design I constructs. **(A)** Fluorescence intensity decay traces measured on H’ labeled with fluorescein and TAMRA (H’_DA) are shown in the absence (red) and presence (black) of IHF. The corresponding donor-only (H’_D) decay traces are also shown in the absence (green) and presence (blue) of IHF. Measurements were done with 5 μM DNA and 5 μM IHF. The instrument response function (gray) is shown for comparison. **(B)** The MEM lifetime distributions obtained from the fluorescence decay traces are shown. The amplitudes of the distributions are normalized to add up to one.

### IHF-H’ complex reveals two dominant DNA conformations: fully-bent (high-FRET) and partially-bent/straight (low-FRET)

We first present lifetime measurements carried out on the H’ DNA substrate, end-labeled with fluorescein and TAMRA (H’ in design I; Table 1), in the presence and absence of IHF. The design of this construct is identical to what we and others have used in previous equilibrium and kinetics studies on IHF-DNA complexes.^60, 68^ All lifetime measurements were done with 5 μM DNA and 5 μM IHF. At 100 mM KCl, the average FRET efficiency in the H’ DNA construct alone was found to be 0.022 ± 0.001, which increased to 0.50 ± 0.03 upon binding of IHF under 1:1 binding conditions. These FRET efficiency values are consistent with steady-state values reported previously for these constructs,^60, 68–69^ and reflect the decrease in the end-to-end distance when H’ DNA is bent into a U-turn in the complex, as illustrated in the crystal structure of the IHF-H’ complex (Figure 1).

MEM analysis on the double-labeled IHF-H’ fluorescence decay traces revealed two dominant lifetime populations for these constructs, with 78 ± 3% of the population in a high FRET state (0.65 ± 0.03), and 22 ± 3% in a low FRET state (0.083 ± 0.002). We note here that the low FRET state is close to what we observe in unbound (straight) DNA, and could arise from (i) a fraction of DNA that is unbound or (ii) a fraction that is nonspecifically bound and that competes with specific binding or (iii) a fraction that is specifically bound but only partially bent, for example if the energetic cost of kinking at two sites is not fully compensated for by stabilizing interactions between protein and bent DNA. Other contributions to this low-FRET state could be from potential artifacts, such as incompletely labeled (or incompletely annealed) DNA, that could result in some fraction of donor-only labeled DNA molecules that are missing an acceptor; partial stacking of the fluorophores at the ends of the DNA that could result in different dye orientations corresponding to stacked or unstacked, and hence two or more distinct FRET states. We ruled out any significant contributions to the observed lifetime distributions from these artifacts, as described in SI Results and SI Figure S2.

### The low-FRET population does not reflect unbound DNA under the 1:1 binding conditions

We first address whether the low FRET population could have contributions from a significant fraction of unbound DNA, even in our 1:1 binding conditions. We note that under buffer and salt conditions (100 mM KCl) identical to those used for these measurements, the *K_d_* of the IHF-H’ complex was previously determined to be 25 pM, from stopped-flow measurements,^60^ and 27 pM, from equilibrium salt-titration measurements.^62^ The binding studies on IHF-H’ presented here are consistent with the previous studies, with an extrapolated value of *K_d_* ≈ 23 pM at 100 mM KCl (SI Figure S4). Therefore, at the protein and DNA concentrations used for the lifetime studies (5 μM each) we expect to have >99.8% of the DNA in complex.

To confirm 1:1 binding conditions for these complexes, we carried out equilibrium titration measurements at a fixed (1 μM) concentration of H’_DA DNA and varying concentrations of IHF, and simultaneously measured both the acceptor ratio (to measure the extent of bent DNA), and donor anisotropy (to measure the extent of protein binding). These data, shown in Figure 3A-B, exhibited behavior that deviated from what is expected for a 1:1 binding isotherm. The acceptor ratio initially increased with increasing protein concentration, as anticipated, and reached a maximum value at [IHF]:[H′] ≈ 1:1; however, the acceptor ratio began to drop with further excess of protein. In contrast, the anisotropy also initially increased with increasing protein concentration, but then continued to increase with further excess of protein beyond [IHF]:[H′] ≈ 1:1. Taken together, we draw the following conclusions from these data: at [IHF]:[H′] ≈ 1:1, each DNA molecule is bound to a single copy of IHF in a specific complex; below this concentration ratio, there is a mixture of unbound and specifically-bound DNA; above this ratio, there is competition between specific and nonspecific binding, with increasing protein concentrations tilting the equilibrium in favor of a nonspecific binding mode in which more than one copy of IHF is bound to a single 35-mer, resulting in higher molecular weight complexes containing less bent DNA (Figure 3C). Competition between specific and nonspecific binding at high IHF to DNA concentrations has been well documented in previous studies.^51, 70–71^

**Figure 3.**
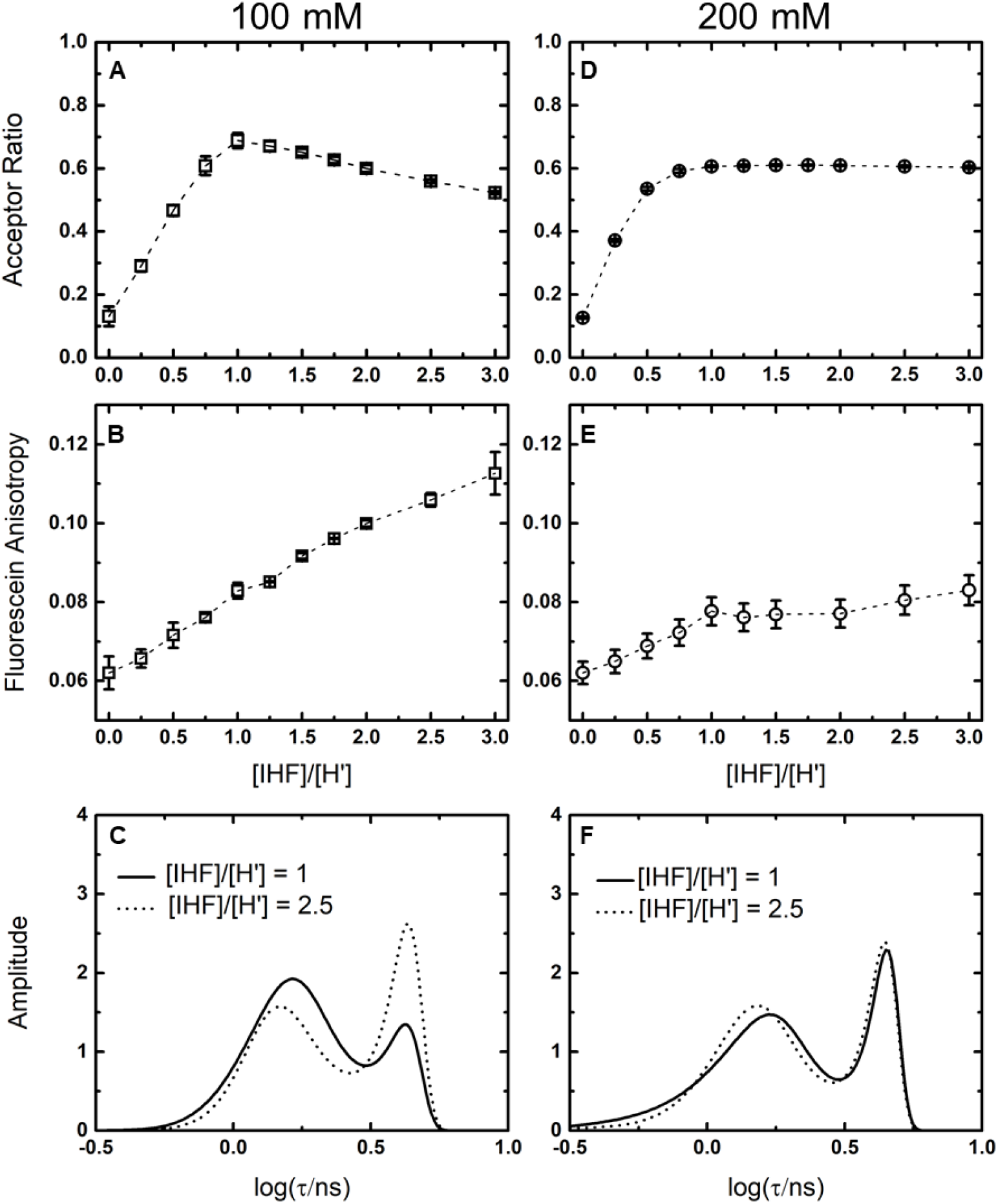
Binding isotherms and MEM distributions for the IHF-H′ complex at 100 and 200 mM KCl. (**A**,**D**) Acceptor ratio measurements and (B,E) anisotropy measurements are shown for 1uM H′_DA and varying concentration of IHF in 100mM KCl (**A**,**B**) and 200mM KCl (**D**,**E**). (**C**,**F**) The MEM lifetime distributions obtained from fluorescence decay traces measured for IHF-H′ are shown for [IHF]/[DNA] = 1 (continuous lines) and [IHF]/[DNA] = 2.5 (dashed lines) in 100 mM KCl (**C**) and 200 mM KCl (**F**). DNA concentrations for the lifetime measurements were 5 μM.

### Increasing the salt concentration diminishes contributions from nonspecific binding

To further examine this competition between specific and nonspecific binding, we performed measurements at a higher salt concentration, 200 mM KCl (Figure 3 and SI Figure S5). Equilibrium studies on the ionic strength dependence of protein-DNA complexes have shown that while higher ionic strength conditions disfavor both binding modes, such conditions are typically more disruptive to nonspecific than to specific binding.^51, 72–75^ In other words, an increase in salt is expected to shift the equilibrium in favor of specific binding, as has indeed been demonstrated by isothermal titration calorimetry (ITC) studies for IHF-H′^51^ as well as for other DNA-bending proteins.^76–79^ This behavior reflects the fact that nonspecific interactions are primarily electrostatic interactions between the protein and the DNA phosphate groups, while specific interactions have significant contributions from hydrogen bonds, van der Waals interactions, and water-mediated interactions that are less affected by ionic strength. Therefore, by increasing the salt concentration from 100 to 200 mM KCl, we anticipated that the contributions from nonspecific binding in our IHF-H′ complexes should diminish, even when IHF is present in excess over DNA.

We repeated our equilibrium titration measurements in 200 mM KCl conditions, with acceptor ratio and anisotropy as probes of the binding process, as before (Figure 3D-E). Indeed, unlike the 100 mM KCl data, the acceptor ratio versus protein concentration profile in 200 mM KCl exhibited the behavior expected for a 1:1 binding isotherm, with the acceptor ratio initially increasing with protein concentration, and then reaching a plateau with protein in excess. The anisotropy data also resembled a 1:1 binding isotherm and did not show any significant evidence for multiple protein binding. Taken together, we infer that at 200 mM KCl nonspecific binding is sufficiently destabilized such that we observe only specific binding at all protein:DNA concentrations.

We now return to the question: what is the origin of the low-FRET component in the lifetime distributions measured on IHF-H′ in 100 mM KCl under 1:1 conditions? Corresponding lifetime measurements for the 1:1 complex at 200mM KCl also showed two populations in the lifetime distributions (Figure 3F), with 66 ± 2% in the high FRET state (*E* ≈ 0.67 ± 0.02) and 34 ± 2% in the low FRET state (*E* ≈ 0.043 ± 0.001). Even with a 2.5-fold increase in the protein concentration (2.5:1 complex) in 200 mM KCl, almost no change was observed in the IHF-H′ lifetime distribution, consistent with our acceptor ratio and anisotropy data at 200mM that showed no evidence for nonspecific binding at excess protein conditions. From these data we conclude that the low-FRET state must be a less-bent DNA conformation of a specifically-bound complex, and which could not be detected in previous steady-state measurements or in crystal structures. The population in this less-bent conformation increased from ˜22% to ˜34% with the increase in salt concentration from 100 to 200 mM KCl.

### Measurements with different placement of FRET labels (design II constructs) implicate the low-FRET state as arising from partial DNA bending

The results presented thus far are insufficient to conclude whether the low-FRET component was from specifically-bound but straight DNA or from one that is partially bent, for example with DNA kinked at only one site or the other. With the FRET labels attached at the ends of the 35-mer H′ substrate (design I in Table 1), the end-to-end distance for a straight piece of DNA is estimated to be ˜119 Å, with near zero FRET efficiency, assuming Forster distance *R*_0_ ≈ 50 Å for this FRET pair.^67, 80^ If we envision a distribution of partially bent conformations with either one side kinked or the other, then the end-to-end distance is estimated to be ˜75 Å, with FRET *E* ≈ 0.08 (not accounting for linker lengths used to attach the labels). These estimates are in good agreement with E ≈ 0.083 ± 0.002 measured for the low-FRET state in IHF-H′ at 100 mM KCl. However, the width in our lifetime distributions makes it difficult to unambiguously separate these low-FRET states from those obtained on straight DNA (Figure 2). To increase the sensitivity of our FRET measurements and more clearly detect FRET changes between straight and partially bent conformations as envisioned above, we altered the labeling strategy to design II (see Table 1), in which the labels are placed closer together. In these constructs, we attached the donor (fluorescein) internally, at a thymine located at position 26 of the top strand (counting from the 5′ end), and attached Atto550 (At) at the 3′ end of the bottom strand. The separation between the labels in these design II constructs is 26 bp (˜88 Å, if unbent).

Fluorescence lifetime decay traces on donor-only H′ DNA in the design II constructs, performed at 100 mM KCl conditions, still exhibited a predominantly single-exponential decay, both in the absence and presence of IHF (Figure 4 and SI Figure S1). The lifetime decay traces for donor-acceptor-labeled H′_DA remained primarily single-exponential, with FRET *E* of 0.113 ± 0.002. In the presence of IHF, the lifetime distributions on the double-labeled constructs showed two distinct populations, reaffirming our results from design I that the DNA in this ensemble sample different conformations, with a high-FRET state observed at 0.52 ± 0.01 and a low-FRET state at 0.29 ± 0.01. Furthermore, in the design II constructs, the FRET efficiency of the low-FRET state of the IHF-DNA complex is well separated from the FRET efficiency of unbound DNA, supporting the conclusion that the low-FRET state corresponds to partially bent DNA. These partially bent complexes could be a mixture of conformations with one side kinked or the other, while the high-FRET state likely corresponds to both DNA sites being kinked and the flanking arms of the DNA held against the sides of the protein, as seen in the crystal structures.

**Figure 4.**
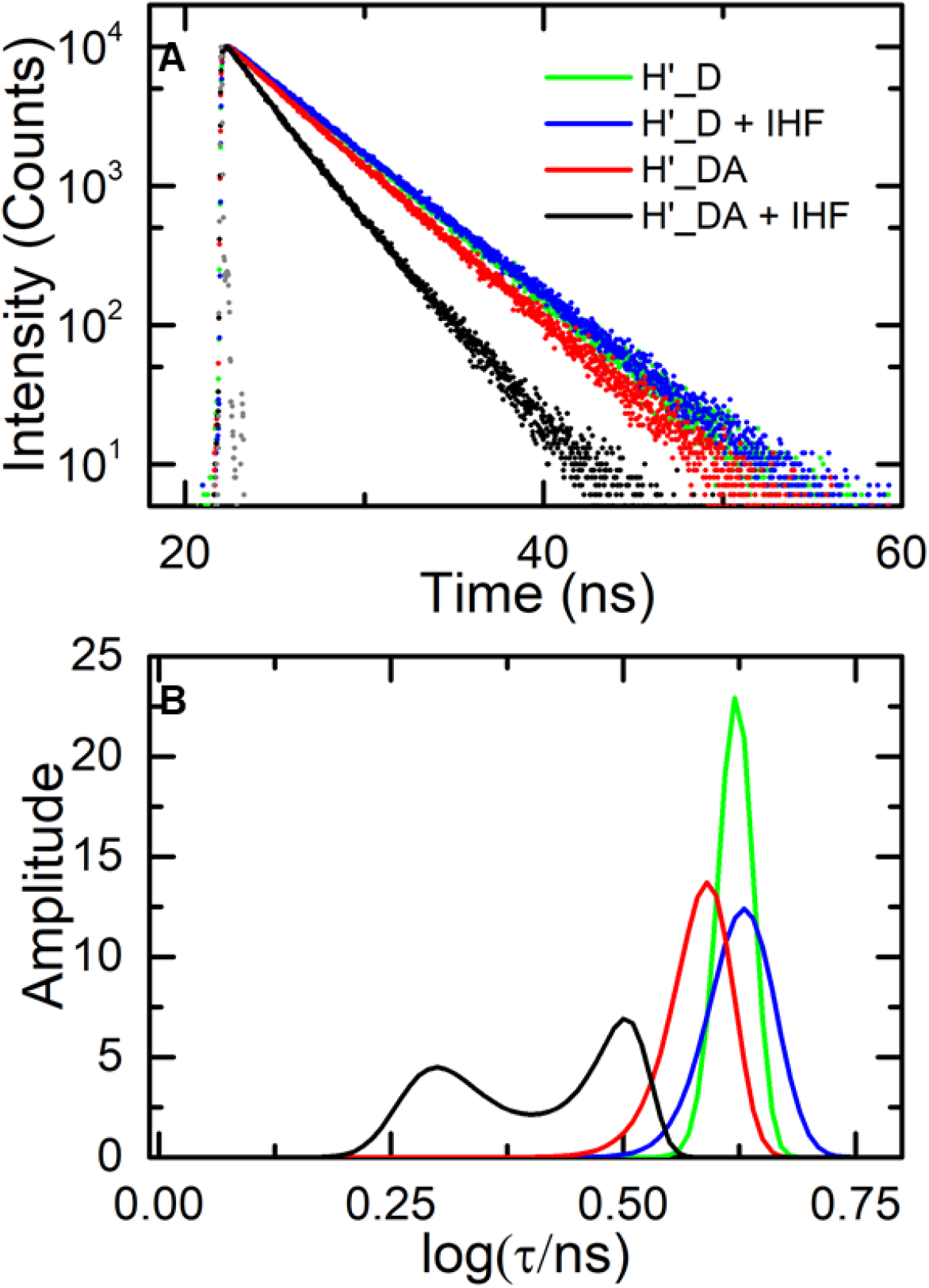
Fluorescence lifetime measurements on IHF-H′ complex in design II constructs. (**A**) Fluorescence intensity decay traces measured on H′ labeled with fluorescein and Atto550 (H′_DA) are shown in the absence (red) and presence (black) of IHF. The corresponding donor-only (H′_D) decay traces are also shown in the absence (green) and presence (blue) of IHF. Measurements were done with 5 μM DNA and 5 μM IHF. The instrument response function (gray) is shown for comparison. (**B**) The MEM lifetime distributions obtained from the fluorescence decay traces are shown. The amplitudes of the distributions are normalized to add up to one.

We note here that the population distribution between the high- and low-FRET states in the design II constructs are split more evenly, with ˜50% in each of these conformations, compared with the approximately 80% versus 20% split observed for the complex in the design I construct under identical conditions. The reason for this population shift could be some steric hindrance between the internally located fluorescein label on the DNA near one of the kink sites that could hamper IHF from keeping the flanking arm of the DNA against its side, although we note that the binding affinity measurements at 300 mM KCl showed no significant difference in the *K_d_* values for IHF-H′ between the two designs (see Table 1). The remaining results presented below are for the design I constructs.

### The H1 site, which lacks the A-tract, favors the less bent state

We next examined the conformational distribution of another specific binding site recognized by IHF, the H1 site on phage lambda DNA. H1 has the same consensus region as H′ (the WATCAAnnnnTTR indicated in gray in Table 1) but differs primarily in the other half of the binding site where it lacks the A-tract that is present in the H′ sequence. Previous stopped-flow measurements revealed a *K_d_* of ˜20 pM for the IHF-H1 complex at 100 mM KCl,^59^ very similar to the *K_d_* from stopped-flow on the IHF-H′ complex.^59–60^ These results indicate that the lack of the A-tract appears to be compensated for by other changes in the H1 sequence compared with H′ so as not to significantly perturb the overall binding affinity. However, the average FRET efficiencies measured for the two end-labeled (design I) constructs in complex with IHF are distinctly different. Our lifetime studies (performed at 100 mM KCl and 20 °C) revealed an average FRET efficiency of 0.25 ± 0.01 for the IHF-H1 complex compared with 0.50 ± 0.03 for IHF-H′ complex (Table 1), consistent with previous steady-state measurements on these complexes using identical constructs and buffer conditions.^59–60, 68^ The decreased FRET efficiency measured in the IHF-H1 complex suggests an inability of IHF to keep the bent arm of the DNA by its side in the absence of the A-tract.

To examine how the presence or absence of the A-tract affects the population distribution of differently bent DNA conformations in the IHF-H1 complex, we analyzed the fluorescence decay traces measured on the end-labeled IHF-H1 complex with the MEM, as before. The lifetime distributions measured for IHF-H1 also show two peaks (Figure 5 and SI Figure S3); however, the DNA in the IHF-H1 complex prefers the low-FRET conformation, with only 36 ± 2% in the high FRET state (*E* ≈ 0.49 ± 0.03) and a larger fraction, 64 ± 2%, in the low FRET conformation (*E* ≈ 0.11 ± 0.01). We assert that these remarkable differences in the conformational distributions of IHF-H′ and IHF-H1 are primarily from the presence and absence of the A-tract, although there are some other changes in the two sequences, especially at the kink sites (see Table 1), that could also be affecting the distributions and the binding affinities. For example, we note that even the high FRET conformation in IHF-H1 (FRET *E* ≈ 0.49) appears to be not as fully bent as in IHF-H′ (FRET *E* ≈ 0.65). Nonetheless, these results showing how the population distribution is affected in the absence of the A-tract highlight the importance of the A-tract and the contacts that it facilitates with IHF to help the protein clamp the bent arm of the DNA on that side. The binding preference of IHF for the A-tract was well noted when the crystal structure was solved, since the A-tract has a unique structure with a narrow minor groove and a high twist which allows it to fit into the protein clamp without significant additional distortions.^9^

**Figure 5.**
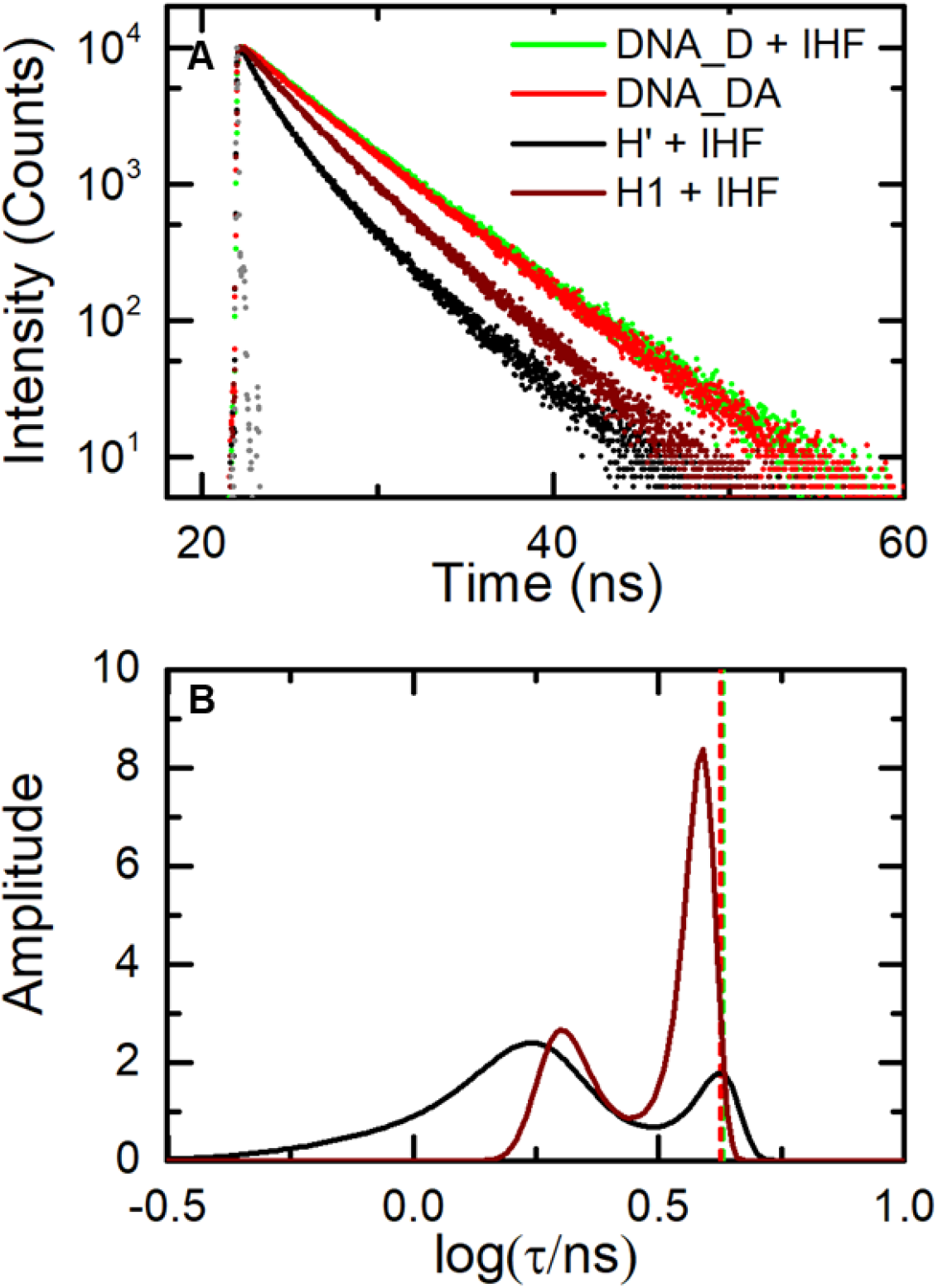
Fluorescence lifetime measurements on IHF-H′ compared with IHF-H1 (design I constructs). **(A)** Fluorescence intensity decay traces are shown for H′_DA (black) and H1_DA (maroon), measured in the presence of IHF. Decay traces on DNA_DA in the absence of IHF (red) and DNA_D in presence of IHF (green) are shown for comparison. Measurements were done with 5 μM DNA and 5 μM IHF. The instrument response function (gray) is shown for comparison. **(B)** The MEM lifetime distributions obtained from the fluorescence decay traces measured for IHF-H′ (black) and IHF-H1 (maroon) are shown. The amplitudes of the distributions are normalized to add up to one. The average lifetime for the DNA_DA in the absence of IHF (red) and DNA_D in presence of IHF (green) are indicated by the vertical dashed lines.

### Insertion of a mismatch at a kink site in H1 on the A-tract side helps recover the U-bent conformation

Next, we inserted a 4 nucleotide bubble (CT/TC mismatch) into the H1 sequence at the kink site on the same side as where the A-tract is in H′ (see H1_CTloop in Table 1). The 4 nt mismatch bubble is expected to enhance the “kinkability” of the H1 sequence on that side and increase the IHF binding affinity, as was shown previously for similar mismatched sequences in the H′ context.^56, 62, 69^ Binding affinity measurements with end-labeled H1_CTloop showed a ˜125-fold decrease in *K_d_* compared with matched H1 (measured at 300 mM KCl; SI Figure S6 and Table 1), a result consistent with previous *K_d_* measurements on H′ sequence with a TT/TT mismatch introduced at the same kink site.^56, 62, 69^ We anticipated that if the shift in population from fully to partially bent DNA in IHF-H1 was due to the missing A-tract, then insertion of the mismatch to lower the energetic penalty for kinking should help compensate for the lack of stabilizing interactions afforded by the A-tract. In other words, we expected to recover some of the lost high FRET state in IHF-H1 in the presence of the mismatch. Our lifetime studies are consistent with this expectation.

First, the average FRET efficiency measured in IHF-H1 increased from 0.25 ± 0.01 in the matched construct to 0.36 ± 0.02 in the mismatched (H1_CTloop) construct (Figure 6 and SI Figure S3). Second, the population in the high-FRET component increased from 36 ± 2% in the matched IHF-H1 to 46 ± 2% in IHF-HI_CTloop. In addition, we observed a shift in the FRET efficiency values for the two populations, with the high FRET state shifted from 0.49 ± 0.03 to 0.54 ± 0.02, and the low-FRET state shifted from 0.12 ± 0.01 to 0.21 ± 0.01, when comparing matched versus mismatched versions of this complex. These shifts in the FRET efficiency levels indicate additional conformational changes in the kinked/straight states of the DNA introduced by the mismatched bubble.

**Figure 6.**
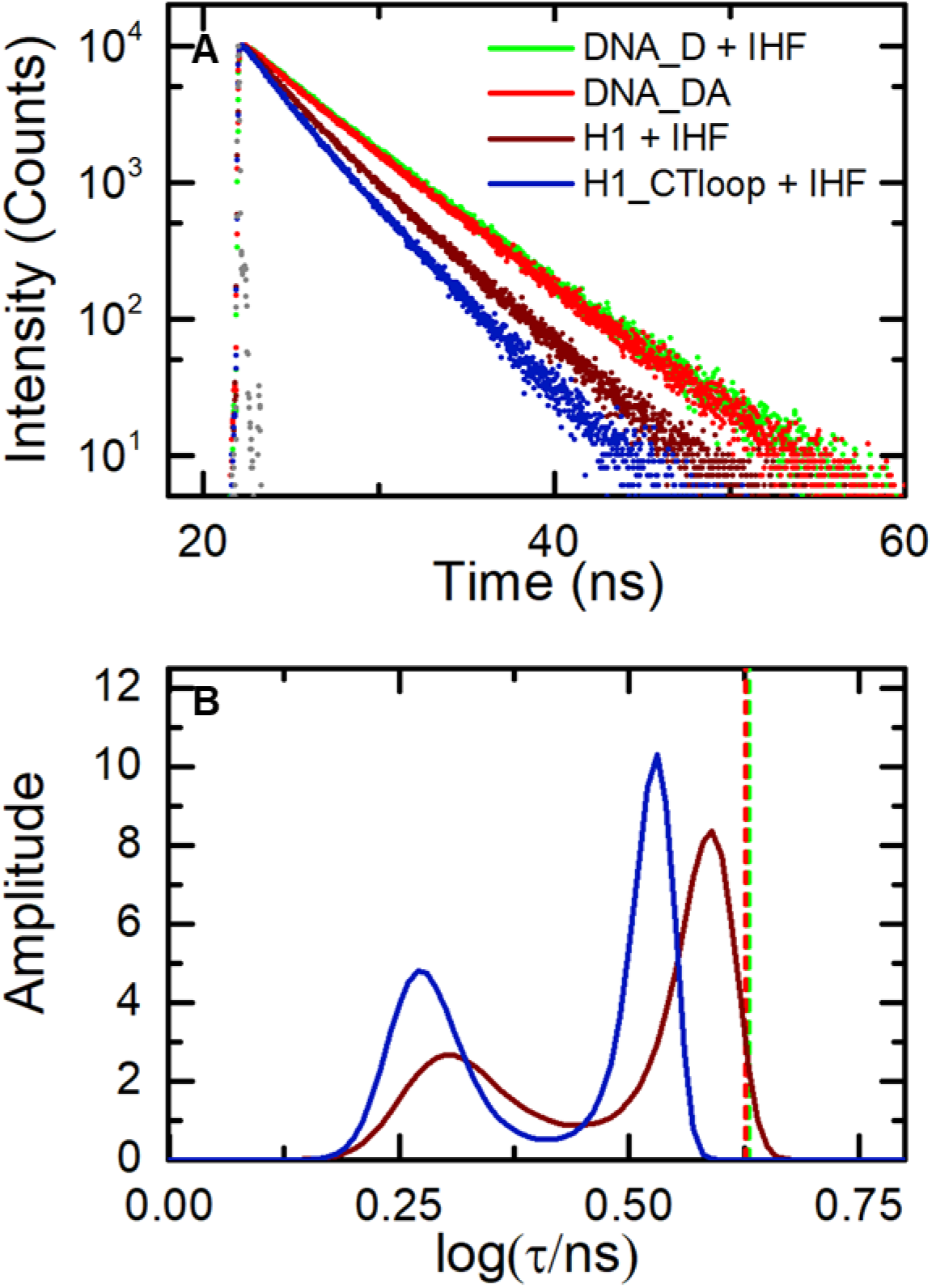
Fluorescence lifetime measurements on IHF-H1 compared for matched and mismatched (design I constructs). **(A)** Fluorescence intensity decay traces measured on H1_DA (maroon) and H1-CTloop_DA (blue), both in the presence of IHF. Decay traces on DNA_DA in the absence of IHF (red) and DNA_D in presence of IHF (green) are shown for comparison. Measurements were done with 5 μM DNA and 5 μM IHF. The instrument response function (gray) is shown for comparison. **(B)** The MEM lifetime distributions obtained from the fluorescence decay traces measured for H1_DA (maroon) and H1_CTloop_DA (blue) are shown. The amplitudes of the distributions are normalized to add up to one. The average lifetime for the DNA_DA in the absence of IHF (red) and DNA_D in presence of IHF (green) are indicated by the vertical dashed lines.

### A destabilizing modification in the TTG consensus region of H′ increases the population in (another) low-FRET state

Next, we examined the effect of sequence modifications in the TTR consensus region on the other flanking arm of the DNA (Figure 1). In this consensus site, a single adenine substitution (TTG→TAG) as shown in the sequence H′44A (Table 1), was found to destabilize the binding affinity of IHF by nearly 100-fold to 250-fold.^57, 62^ The crystal structure for the IHF-H′44A complex revealed that this single nucleotide substitution inhibited the twisting of the DNA that was needed to form a network of salt bridges with IHF that stabilized the bent DNA conformation against that side of the protein.^57^ However, the crystal structure of IHF-H′44A did reveal a fully bent conformation very similar to that of IHF-H′. In contrast, steady-state FRET measurements on the IHF-H′44A complex at 20 °C in solution yielded a smaller average FRET efficiency than IHF-H′ for the design I constructs under identical binding conditions,^62^ indicating some degree of “floppiness” in the bent conformations of IHF-H′44A.

To examine how this T→A mutation in the TTG region affects the populations in the fully versus partially bent conformations of the IHF-bound specific complexes, we carried out lifetime studies on the end-labeled (design I) IHF-H′44A complex (Figure 7 and SI Figure S3). We still observe two populations as for the IHF-H′ complex, with a high FRET state at *E* ≈ 0.59 ± 0.03 and a low FRET state at *E* ≈ 0.042 ± 0.001, but with a significant decrease in the high FRET population of 36 ± 1% compared with 78% in IHF-H1. These results demonstrate the importance of the TTG consensus region in keeping that side of the DNA clamped against the protein and further show that neither of the flanking DNA arms are held rigidly in place. Altogether, these results suggest at least three conformations that co-exist in solution for a specially bound protein: a fully bent complex and two partially bent complexes with one arm straight or the other. The results from IHF-H′44A suggest that the population in the second partially bent complex that affects the TTG side of the complex increases by 42% when that side is compromised.

**Figure 7.**
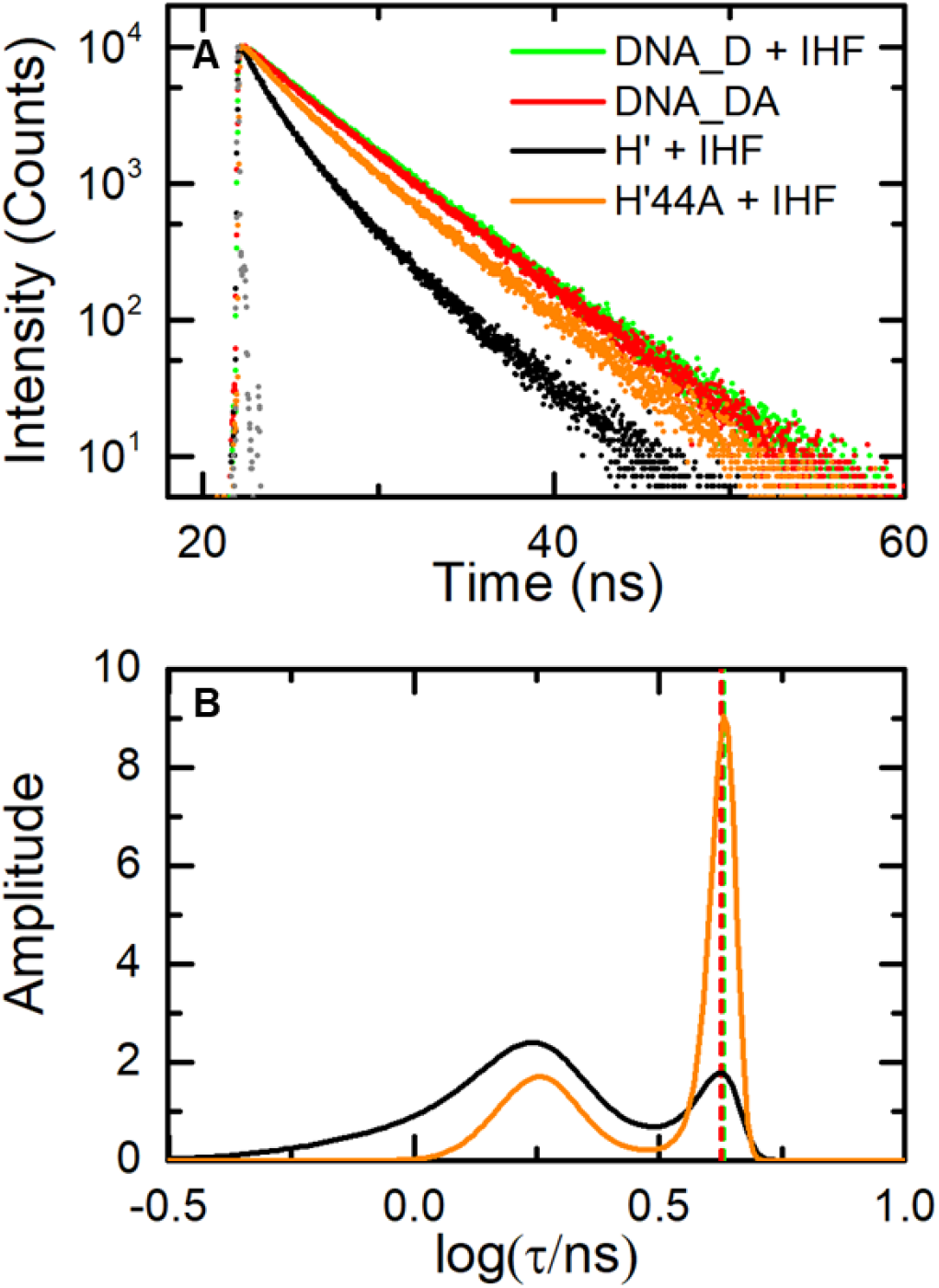
Fluorescence lifetime measurements on IHF-H′ compared with IHF-H′44A (design I constructs). **(A)** Fluorescence intensity decay traces measured on H′_DA (black) and H′44A_DA (orange), both in the presence of IHF. Decay traces on DNA_DA in the absence of IHF (red) and DNA_D in presence of IHF (green) are shown for comparison. Measurements were done with 5 μM DNA and 5 μM IHF. The instrument response function (gray) is shown for comparison. **(B)** The MEM lifetime distributions obtained from fluorescence decay traces measured for H′_DA (black) and H′44A_DA (orange) in the presence of IHF are shown. The amplitudes of the distributions are normalized to add up to one. The average lifetime for the DNA_DA in the absence of IHF (red) and DNA_D in presence of IHF (green) are indicated by the vertical dashed lines.

## DISCUSSION

IHF is a small protein belonging to a class of nucleoid-associated DNA-bending proteins. Apart from its nonspecific biological function in condensing the bacterial nucleoid, it also binds in a sequence-specific manner and serves as an architectural factor in many cellular activities such as site-specific recombination, DNA replication and transcription.^81–82^ The ability of IHF to bend the DNA containing its specific site into a U-turn by wrapping ˜35 bp DNA around three sides of the protein has earned it the moniker of the “master bender”.^83^ Remarkably, IHF accomplishes this feat with almost no direct interactions between the protein residues and specific bases, and has thus become an excellent model system for studies of sequence-dependent DNA shape and deformability that underpins binding site recognition by indirect readout.^71, 84–86^

The sharp DNA bends induced by IHF allow for FRET measurements to be sensitive reporters of the extent of DNA bending. Previous studies took advantage of time-resolved FRET to investigate DNA-bending kinetics in IHF-DNA complexes,^60, 68^ which demonstrated the stepwise binding-then-bending mechanism for site recognition by IHF.^87^ Further kinetics studies resolved a two-step “interrogation-then-recognition” process: nonspecific interrogation on ˜100-500 μs timescale prior to recognition on 1-10 ms timescale.^69^ However, these “ensemble” approaches could only provide an average picture of the dynamics along the reaction trajectory. What remained elusive was whether multiple conformations of the complex could co-exist in solution, as was recently shown for damaged DNA specifically bound to NER damage-recognition protein XPC/Rad4.^43^ Here, we investigated the distribution of bent conformations in IHF-DNA complexes with varying DNA sequence composition and binding affinities. We utilized fluorescence lifetime decay measurements and the maximum entropy methods (MEM) to obtain FRET distributions that enabled us to visualize and quantitatively characterize the heterogeneity of bent conformations.

Like many DNA-bending proteins that kink DNA at localized sites, IHF concentrates the U-shaped bend in two sharp kinks separated by 9 bp. At the kink sites, a single base-step is unstacked and opened towards the minor groove of the DNA, and stabilized by intercalation of conserved proline residues on the β-arms of the protein that wrap around the DNA. The flanking sides of the DNA on the outer sides of the kinks are held against the body of the protein through a myriad of specific and nonspecific interactions. Notably, the consensus DNA-binding motif consists of only a 6-bp stretch (WATCAR) in between the kink sites and another 3-bp stretch (TTR) in the flanking DNA, making it all the more remarkable that IHF is able to overcome the energy penalty needed to severely deform the DNA at its preferred sites and bind with affinities that can exceed 10^3^-10^4^-fold compared with random sequences.^88–91^

How accurate is this picture of bent DNA rigidly held against the protein, as implied by the static crystal structure of the IHF-H′ complex? Incorporation into a crystal can “freeze out” macromolecular dynamics and will tend to select a single conformation from the ensemble that may exist in solution. The results reported here demonstrate that IHF does in fact experience some difficulty in keeping the bent arms of the DNA at its side. For the IHF-H′ complex in 100 mM KCl, the ensemble of bent conformations appears as two populations, as measured by the FRET distributions on labeled DNA constructs. For the end-labeled (design I) constructs, the population in the fully-bent high-FRET state (*E* ≈0.65) is found to be 78%, with the remaining 22% in a low-FRET state (*E* ≈ 0.08). Although measurements on the design I constructs could not readily distinguish between partially bent or unbent DNA, FRET measurements with design II constructs, where the FRET labels are placed closer together, establish that the low-FRET state is not from unbent (straight) DNA. The low-FRET state is attributed to a partially bent but still specific complex, as clarified from measurements at 200 mM KCl, where nonspecific binding is disfavored, and yet the amplitude of the low-FRET state increases. From these and further results discussed below, we attribute the low-FRET state to an ensemble of specifically-bound conformations with either one or the other kink site unkinked (Figure 8). The free energy of the fully bent IHF-H′ conformation is estimated to be 1.3 *k_B_T* lower than the partially bent ensemble in 100 mM KCl and 0.7 *k_B_T* lower in 200 mM KCl (Table 2). Notably, the FRET efficiency levels observed in the fully-bent conformation are very similar in both ionic conditions.

**Figure 8.**
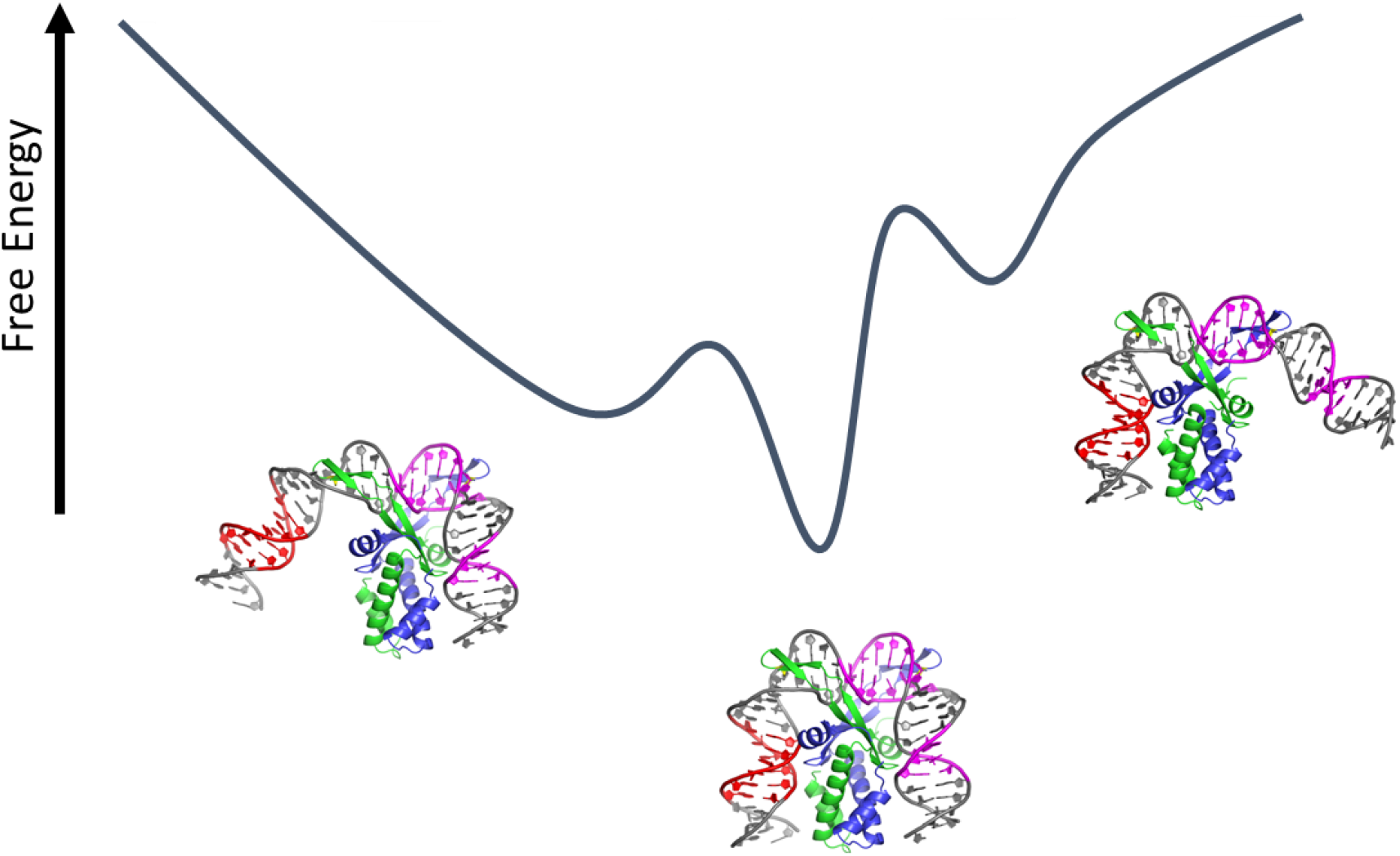
Schematic representation of the free energy landscape of the specific IHF-DNA complex, with multiple conformations accessible in solution. These conformations include the most stable complex, with two DNA sites kinked, as well as conformations with one or the other side unkinked. The partially bent conformations shown are models of the low-FRET population and not based on real structural data.

**Table 2:** Lifetime Measurements on IHF-DNA Complexes

| Construct | $\langle \tau_1 \rangle$ (in ns) | $\langle E_1 \rangle$ | $\alpha_1$ | $\langle \tau_2 \rangle$ (in ns) | $\langle E_2 \rangle$ | $\alpha_2$ | $\Delta G/k_B T = -\ln\left(\frac{\alpha_1}{\alpha_2}\right)$ |
| --- | --- | --- | --- | --- | --- | --- | --- |
| IHF-H' (design I) | $1.52 \pm 0.06^a$<br>($1.41 \pm 0.03$)<sup>b</sup> | $0.65 \pm 0.03$<br>($0.67 \pm 0.02$) | $78 \pm 3\%$<br>($66 \pm 2\%$) | $3.96 \pm 0.01$<br>($4.13 \pm 0.02$) | $0.083 \pm 0.002$<br>($0.043 \pm 0.001$) | $22 \pm 3\%$<br>($34 \pm 2\%$) | $-1.2 \pm 0.2$<br>($-0.66 \pm 0.04$) |
| IHF-H' (design II) | $2.07 \pm 0.01$ | $0.52 \pm 0.01$ | $50 \pm 2\%$ | $3.05 \pm 0.01$ | $0.28 \pm 0.01$ | $50 \pm 2\%$ | $\approx 0$ |
| IHF-H1 (design I) | $2.19 \pm 0.14$ | $0.49 \pm 0.03$ | $36 \pm 2\%$ | $3.80 \pm 0.12$ | $0.12 \pm 0.01$ | $64 \pm 2\%$ | $0.58 \pm 0.04$ |
| IHF-H'44A (design I) | $1.75 \pm 0.07$ | $0.59 \pm 0.03$ | $36 \pm 1\%$ | $4.14 \pm 0.05$ | $0.042 \pm 0.001$ | $64 \pm 1\%$ | $0.58 \pm 0.02$ |
| IHF-H1_CTloop (design I) | $1.99 \pm 0.07$ | $0.54 \pm 0.02$ | $46 \pm 2\%$ | $3.42 \pm 0.18$ | $0.21 \pm 0.01$ | $54 \pm 2\%$ | $0.16 \pm 0.01$ |

<sup>a</sup> Measurements were done in 100 mM KCl
<sup>b</sup> Values in parenthesis are for measurements done in 200 mM KCl

This tug-o-war between protein-induced DNA kinking and the propensity of DNA to retain an unkinked conformation is further illustrated by measurements with the H1 binding site that is missing an A-tract found in the H′ site, which is known to help stabilize the bent DNA conformation in the IHF-H′ complex.^9, 50^ FRET distributions on IHF-H1 revealed significantly less population (36%) in the high-FRET conformation, indicating that the free energy of the fully-bent IHF-H1 conformation is 0.58 *k_B_T* higher than the partially bent conformations. Furthermore, the protein-DNA interactions in the fully bent IHF-H1 conformation (FRET *E* ≈ 0.49) appear to be looser than that in the fully-bent IHF-H′ (FRET *E* ≈ 0.65). Insertion of 2-bp mismatches at one of the kink sites in H1 close to the side that is missing the A-tract (H1_CTloop in Table 1), designed to make the DNA more “kinkable”, compensates to some extent for the loss of the A-tract; the fully-bent complex in IHF-H1_CTloop is still less favored but now only by 0.16 *k_B_T* (Table 2). Thus, the A-tract helps to maintain a tight fit in the complex and its loss results in significant unkinking on that side. These results are in accord with previous hydroxyl radical footprinting studies that showed less protection in that patch in sequences that were missing the A-tract.^46, 84^

Another important and highly conserved feature common to all known binding sites of IHF is the TTR consensus region on the other flanking side of the DNA. Previous studies identified that a T→A switch at the center position of the TTR element (H′44A) resulted in >100-fold decrease in binding affinity for IHF.^57, 62, 92^ Comparison of IHF-H′ and IHF-H′44A structures showed that the ability of the TTR site to adopt an unusually highly twisted conformation at the Y-R step when bound to IHF was necessary to facilitate stabilizing salt bridges between key residues of IHF.^57^ These studies exemplified how sequence-dependent DNA deformability was critical to the recognition of that consensus site.

The crystal structures of IHF-H′ and IHF-H′44A were otherwise very similar, with approximately the same overall bend in the DNA observed in both structures.^57^ In contrast, previous steady-state FRET studies that monitored the average end-to-end distance already revealed a less bent conformation for IHF-H′44A in solution, with FRET efficiency levels nearly half of what was observed in IHF-H′.^62^ Our lifetime measurements directly show that indeed the fraction in the fully-bent conformation of IHF-H′44A is only 36%, indicating a penalty (ΔΔ*G*) of nearly 1.9 *k_B_T* between the bent and unbent states that is attributable to the loss of the salt bridge interactions at the TTR site. Furthermore, the bent state itself is less stably bent, as indicated by a slightly lower FRET efficiency of 0.59 in the high-FRET state of IHF-H′44A compared with 0.65 in IHF-H′.

It is informative to compare the conformational distributions measured here for the specific IHF-DNA complexes with the conformational distributions for the structurally similar but nonspecific HU protein observed in AFM studies.^25^ HU is known to bind in a sequence-independent manner to DNA with *K_d_* values that range from 200 nM to 2.5 μM,^24, 93^ and with much higher affinities to distorted DNA.^55, 93–94^ Single-molecule micromanipulation studies of HU-bound DNA showed that at low HU/high monovalent salt concentrations, HU dimers induce very flexible bends that result in DNA compaction and a dramatic decrease in the apparent persistence length of DNA compared with bare DNA.^25, 95^ AFM studies under similar conditions revealed a very broad range of bend angles in the DNA at the sites where HU was bound,^25^ with a nearly uniform distribution of angles from 0° (unbent) to 180° (bent into a U-shape). Together, these studies revealed a highly compliant and very flexible HU-DNA complex.

Similar conclusions were drawn from force-extension measurements on long DNA with IHF bound nonspecifically, that also showed enhanced apparent DNA flexibility with the bound proteins.^91, 96^ AFM studies with specific IHF-DNA complexes revealed single broad distributions peaked at bending angles of ˜120-130°, with a range that covered bending angles from ˜80° to ˜160°.^27, 33^ These AFM studies were done with other IHF-DNA complexes; we are unaware of similar studies with IHF-H′. Notably, the range of bent conformations observed in AFM covers what we expect for the low-FRET state (with one site kinked) and for the high-FRET state (with both sites kinked).

However, our studies indicate a somewhat “brittle” complex for IHF bound to its specific site. Rather than describing a broad and continuous range of bent conformations as seen in AFM images of HU-DNA and IHF-DNA complexes, our data support two or likely three preferred conformations, with the populations among these distinct valleys in the free energy landscape modulated by the DNA sequence.

In a previous smFRET study on IHF-H′, using a 55-bp DNA construct containing the H′ site and end-labeled with a FRET pair, a bimodal FRET distribution was indeed observed, with ˜85% in a high-FRET conformation consistent with the crystal structure and ˜15% population in what appeared as a “zero-FRET” conformation. Remarkably, a very similar bimodal distribution was also observed in complexes of HU bound to a 55-mer with two TT mismatches 9 bp apart, but not with HU bound to the 55-mer H′ construct, which revealed a broad, feature-less distribution reflecting less severely bent conformations. HU has been shown to bind with very high affinity (Kd in the 4-10 pM range) to DNA substrates with mismatches spaced 9 bp apart and it is not unexpected that HU can induce U-bends in these high affinity sequences similar to IHF-H′. The authors of this study interpreted the zero-FRET component as arising from nonspecifically bound proteins. Indeed, it is not evident that these smFRET measurements could discern distinct populations within the specific complex, if those conformations interconverted on timescales faster than (or comparable to) the 1 ms binning times of this smFRET study.^97^ Kinetics measurements on IHF-DNA complexes indicate that DNA bending/unbending dynamics within the complex are indeed fast, on micro-to-millisecond timescale.^60, 62, 68–69^

Another smFRET study, designed to examine rigid versus flexible kinks for another nonspecific DNA-bending protein from the eukaryotic family of HMG box proteins, concluded that the ˜60° kinks induced by that protein appeared to be rigid kinks, with the apparent enhancement of DNA flexibility induced by these proteins attributed to binding/unbinding of the protein to induce random and transient kinks.^98^ Again, as the authors of that study noted, the ˜30 ms binning time of their smFRET measurements could have averaged out any dynamic flexibility. Further studies, including measurements of the kind reported here, are needed to carefully flesh out the dynamics and distributions of these ubiquitous DNA benders.^99^

Our observation of partially bent specific conformations in IHF-DNA complexes also provide structural insights into the underlying mechanism for the “facilitated dissociation” observed for several protein-DNA complexes, whereby dissociation rates of these complexes are found to depend on the protein concentrations.^100–104^ As postulated by others,^103, 105–106^ for dimeric DNA-binding proteins, the release of a monomer from a half-site is a plausible mechanism to generate a partially bound intermediate, thereby making room for another protein to bind and eventually displace the first protein. Our measurements provide direct evidence for analogous partially bound structures in IHF-DNA complexes, in which the unkinked DNA arm could interact with a second IHF dimer before the first one fully dissociates.

Finally, we propose that the details of IHF binding sites have evolved to fit their biological roles (Figure 9). In particular, the A-tract that clearly helps to keep the DNA bent in the U-shape is not conserved across the many known IHF sites, H1 being a case in point. Binding of IHF to both the H′ and H1 sites is required for integration of phage lambda into the *E. coli* chromosome to establish lysogeny. The H′ and H1 sites are found within the “attP” region of phage lambda, which is bound synergistically by 3 copies of IHF and 4 copies of lambda integrase to form a large complex (termed an intasome) that then binds the bacterial insertion site (“attB”) and catalyzes a site-specific recombination reaction between attP and attB that results in the integration of the phage DNA into that of the host. Modeling of this intasome shows that, while the IHF-H′ complex might need to flex slightly to allow synergetic binding of two domains of integrase to DNA sequences flanking it, it can remain static throughout the integration reaction.^58^ However, in the fully bent form of the IHF-H1 complex, the flanking DNA and the copy of integrase bound to it block incorporation of the bacterial attB DNA segment. Significant flexing of the H1-induced DNA bend as shown in Figure 9 is therefore required for its biological function.

**Figure 9.**
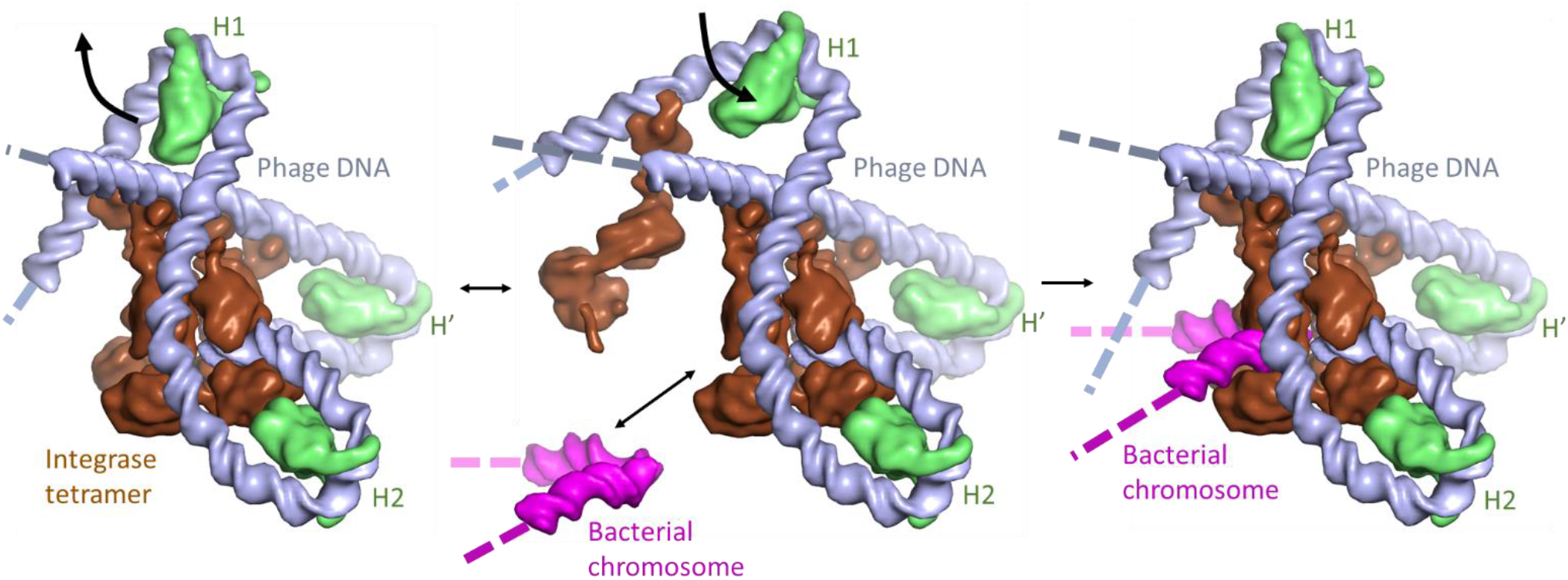
Flexible DNA bending at the H1 site may facilitate assembly of the phage lambda integration complex. Proteins and DNA segments in the model are shown as smoothed surfaces. The “intasome” assembles on phage DNA (“attP”; gray), with 3 copies of IHF (green) bending the DNA such that the integrase tetramer (brown) can bridge multiple DNA sites. Transient flapping of the IHF-induced bend at the H1 site (center panel) allows insertion of the bacterial integration site DNA (“attB”; magenta) into the complex, which is then trapped by closure of the H1 bend. Note that in these images the H1 binding site is oriented such that the (missing) A-tract side is on the right, i.e. orientation of IHF relative to the DNA at the H1 site is flipped 180° from that shown in Figure 1. This figure was made using the PyMol molecular graphics system, version 2.0, Schrodinger, LLC.

## CONCLUSION

The present study showcases the power of combining fluorescence lifetime measurements with MEM analysis for investigating conformational flexibility in protein-DNA complexes and establishes conclusively that IHF bound to its specific sites samples two or more distinct conformations. These conformations include a fully-bent conformation such as that observed in the crystal structures of IHF-DNA and competing conformations in which very likely either one kink site or the other is unkinked. The equilibrium distribution between these different conformations depends sensitively on DNA sequence, especially the A-tract on one side of the U-bend and the TTR consensus site on the other side. The “kinkability” at the kink sites also has a measurable effect on the distribution. Further studies of this nature would be very useful in characterizing DNA sequences that render DNA highly “kinkable” and data such as these could be used to further refine models for sequence-dependent DNA deformability and protein-DNA interactions needed to stabilize distorted DNA conformations in complex with DNA-bending proteins.

## Supporting Information

SI Methods 1.1-1.4; SI Results; SI Figures S1-S6.

## Acknowledgments

We thank Ying Z. Pigli for the purification of IHF proteins and Greg Van Duyne for kindly providing the coordinates for the integrative intasome model. This work was funded by National Science Foundation (NSF) grants MCB-1715649 (to A.A. and P.A.R.) and an LAS Award for Faculty in the Sciences from the University of Illinois at Chicago (to A.A.) This work was also supported by the Intramural Research Program of the National Institutes of Health (NIH, CIT).

---

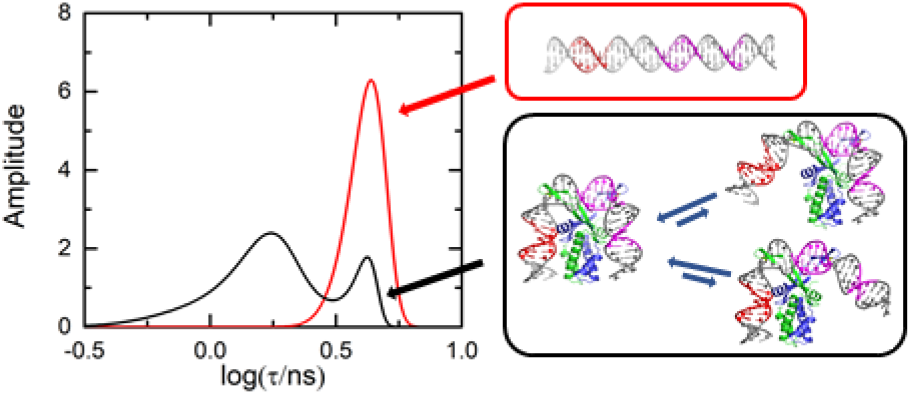
TOC Figure.

